# Whole-genome sequencing of an advanced case of small-cell gallbladder neuroendocrine carcinoma

**DOI:** 10.1101/052316

**Authors:** Maolan Li, Fatao Liu, Yijian Zhang, Xiangsong Wu, Wenguang Wu, Xu-An Wang, Shuai Zhao, Shibo Liu, Haibin Liang, Fei Zhang, Yuan Gao, Shanshan Xiang, Huaifeng Li, Wei Lu, Hao Weng, Jiasheng Mu, Yijun Shu, Runfa Bao, Lin Jiang, Yunping Hu, Wei Gong, Yun Zhang, Tieliang Ma, Kai Zhang, Yun Liu, Yingbin Liu

**Affiliations:** Department of General Surgery, Xinhua Hospital Affiliated to Shanghai Jiao Tong University School of Medicine, Shanghai, 200092, China.; Institute of Biliary Tract Disease, Shanghai Jiao Tong University School of Medicine, Shanghai, 200092, China.; Dapartment of Hepatobiliary and Laparoscopic Surgrey, the Affiliated Yixing Hospital of Jiangsu University, Yixing, 214200, China.; Institutes of Biomedical Sciences, Fudan University, Shanghai, 200433, China.

**Author notes:** These authors contributed equally to this work (Maolan Li, Fatao Liu, Yijian Zhang). These authors jointly directed this work. Correspondence should be addressed to Y.B.L., Y.L. or K.Z.

## Abstract

The majority of gallbladder cancer cases are discovered at later stages, which frequently leads to poor prognoses. Small-cell gallbladder neuroendocrine carcinoma (GB-SCNEC) is a relatively rare histological type of gallbladder cancer, and its survival rate is exceptionally low because of its greater malignant potential. In addition, the genomic landscape of GB-SCNEC is rarely considered in treatment decisions. We performed whole-genome sequencing on an advanced case of GB-SCNEC. By analyzing the whole-genome sequencing data of the primary cancer tissue (76.29X coverage), lymphatic metastatic cancer tissue (73.92X coverage) and matched non-cancerous tissue (35.73X coverage), we identified approximately 900 high-quality somatic single nucleotide variants (SNVs), 109 of which were shared by both the primary and metastatic tumor tissues. Somatic non-synonymous coding variations with damaging impact in HMCN1 and CDH10 were observed in both the primary and metastatic tissue specimens. A pathway analysis of the genes mapped to the SNVs revealed gene enrichment associated with axon guidance, ERBB signaling, sulfur metabolism and calcium signaling. Furthermore, we identified 20 chromosomal rearrangements that included 11 deletions, 4 tandem duplications and 5 inversions that mapped to known genes. Two gene fusions, NCAM2-SGCZ and BTG3-CCDC40 were also discovered and validated by Sanger sequencing. Additionally, we identified genome-wide copy number variations and microsatellite instability. In this study, we identified novel biological markers of GB-SCNEC that may serve as valuable prognostic factors or indicators of treatment response in patients with GB-SCNEC with lymphatic metastasis.

## Introduction

Gallbladder cancer is the most common aggressive tumor of the biliary tract and accounts for 80-95% of these types of cancers worldwide(Lai and Lau 2008). Because symptoms are frequently indiscernible in the early stages, patients are often diagnosed at more advanced stages of the disease. As a result, the median 5-year survival rate associated with gallbladder cancer remains low(Zhu et al. 2010). Adenocarcinomas account for most gallbladder malignancies (90%), which include the less common squamous, adenosquamous, and neuroendocrine carcinoma (NEC) histologic subtypes. NEC commonly arises in the gastrointestinal tract and the bronchi but is rarely observed in the gallbladder(Yao et al. 2008). Small-cell neuroendocrine carcinoma (GB-SCNEC) of the gallbladder is even less common and has an incidence of approximately 0.5% of all gallbladder cancers(Henson et al. 1992) and 0.2% of all gastrointestinal carcinomas according to Surveillance, Epidemiology and End Results (SEER) data(Jun et al. 2006). The symptomatology of GB-SCNEC is nonspecific, and the diagnosis of GB-SCNEC is often based on the detection of cholecystolithiasis or polyps by cholecystectomy. Women and patients with cholelithiasis are considered high-risk populations for GB-SCNEC, and approximately 75% of patients from these populations present with concomitant hepatic, pulmonary, peritoneal and lymph node metastases. Because of the rarity of this disease, standards of care for treatment have not been clearly defined, and effective treatment options are currently limited to complete surgical resection with negative margins(El Fattach et al. 2015). Although GB-SCNEC cases represent a relatively small proportion of the total gallbladder cancer cases, the survival rate for neuroendocrine carcinomas is much less that the overall rate for gallbladder cancer because of its malignant potential and late-stage diagnosis. Thus, novel insights into the molecular and genetic aberrations specific to GB-SCNEC might facilitate the development of more efficient methods for the prevention, early diagnosis and curative treatment of this disease.

Gene mutations associated with gallbladder cancers are typically somatic in nature rather than inherited (2015). Mutations and overexpression of the p53 gene are the most common genetic changes observed in human cancers(Itoi et al. 1997), and p53 mutations are observed in up to 70% of the gallbladder cancer cases evaluated in various studies(Saetta 2006). In a previous study, we observed a significant frequency of non-silent p53 mutations in gallbladder cancer (47.1%)(Li et al. 2014). K-ras is another well-studied gene implicated in gallbladder cancer. Recent investigations have shown that K-ras mutation rates in various cancers range from 10 to 67%(Saetta 2006). We previously reported that a substantial proportion of K-ras mutations in gallbladder cancer are non-silent (7.8%), which is similar to p53(Li et al. 2014). The ERBB signaling pathway has the most well-established association with gallbladder carcinogenesis. Mutations in the components of the ERBB signaling pathway, including EGFR, ERBB2, ERBB3 and ERBB4, are the most frequently observed mutations in gallbladder cancer and affect approximately 37% of all gallbladder tumors(Li et al. 2014). In a recent study, Miyahara N demonstrated that Muc4 activation of ERBB2 signaling contributed to gallbladder carcinogenesis(Miyahara et al. 2014). Microsatellite instability (MSI) is another notable genetic factor associated with gallbladder cancer, and the frequency of MSI ranges from 3% to 35%(Saetta 2006).

Whole-genome sequencing (WGS) is considered a more powerful method for identifying putative disease-causing mutations in exonic regions compared with whole-exome sequencing (WES)(Belkadi et al. 2015); however, WGS studies in gallbladder cancer and metastatic gallbladder cancer have not been reported. The present study is the first to report the application of WGS technology to study genome-wide single nucleotide variants (SNVs), structural variations, copy number variations (CNVs) and microsatellite stability in a patient with GB-SCNEC. We also evaluated genomic differences between primary and metastatic GB-SCNEC tissues. Together, our findings might provide insights into genome-wide aberrations that mediate gallbladder carcinogenesis and metastasis.

## Materials and Methods

### 1. Clinicopathological characteristics of the patient

A 56-year-old man was hospitalized with a 1-month history of pain in the right hypochodrium. The physical examination revealed no abnormalities. On admission, tumor marker levels were within the normal range. An abdomen CT with and without contrast revealed markedly abnormal gallbladder morphology with an enlargement at the neck region measuring 4.7 × 4.5 cm. The porta hepatis lymph nodes were abnormally enlarged, and stones were not visible in the gallbladder mass (**Figure 1A**, **1B**). No neoadjuvant or adjuvant chemotherapy was administered prior to surgery. The surgical procedure (which included cholecystectomy as well as extensive surgical resections, including segment 4 of the liver and regional lymph node clearances) was performed in July 2014. After the surgical resections, histopathological analyses revealed that the tumor was composed of monomorphic cells containing small round nuclei and eosinophilic cytoplasm. The cells were organized in small nodular, trabecular or acinar structures surrounded by highly vascularized stroma with a high mitotic index (**Figure 1C**, **1D**). Immunohistochemistry assays revealed that the cells in both the tumor and the resected lymph nodes were negative for chromogranin A but positive for synaptophysin, NSE and Ki 67 (80% of cells) (**Figure 1E-1L**). These findings were consistent with a diagnosis of GB-SCNEC with lymph node involvement. All of the surgical margins were negative, and a pathological R0 resection was achieved. The patient later received systemic chemotherapy with VP-16 and cisplatin and died 17 months after the operation.

**Figure 1.**
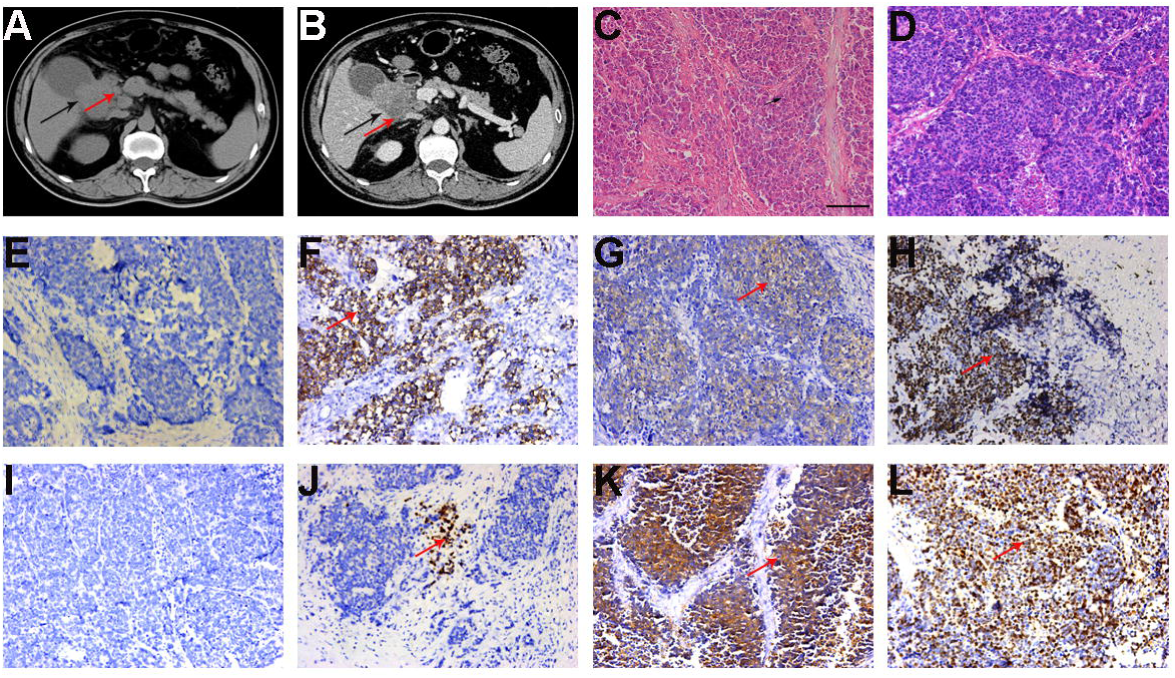
Clinical data of the patient with GB-SCNEC. (A, B) MDCT of the abdomen in the axial plane after the intravenous administration of iodinated contrast material revealed a heterogeneous mass (black arrow) in the neck of the gallbladder in association with large heterogeneous lymphadenopathies (red arrow) in the porta hepatis space. (C, D) Microscopic features of primary gallbladder cancer (C) and metastatic lymph node tissues (D) observed using hematoxylin and eosin (H&E) staining. (E-L) Immunohistochemistry analysis revealed that the tumor cells were negative for chromogranin A but positive (red arrow) for synaptophysin, NSE and Ki67 (80% of cells) in the primary gallbladder cancer (E-H) and metastatic lymph node tissues (I-L).

### 2. Whole-genome sequencing

Genomic DNA was extracted from the tissue specimens using the QIAamp DNA kit (Qiagen). The libraries were prepared using the protocols provided by Illumina. Briefly, 1 μg of DNA was sheared into small fragments (approximately 200-300 bp) using a Covaris S220 system. DNA fragments were end repaired and 3’ polyadenylated. Adaptors with barcode sequences were ligated to both ends of the DNA fragments. E-Gel was used to select the target-size DNA fragments. Then, the tissues underwent 10 cycles of PCR amplification, and the resulting PCR products were purified. Validated DNA libraries were sequenced using the Illumina Sequencing System (HiSeq Series).

### 3. Whole-genome sequencing data processing and mutation identification

Read pairs (FASTQ data) generated from the sequencing system were trimmed and filtered using Trimmomatic(Bolger et al. 2014). The resulting reads were mapped to the human reference genome hg19 downloaded from the UCSC Genome Browser using BWA-MEM (http://arxiv.org/abs/1303.3997)) in the SpeedSeq framework(Chiang et al. 2015). Duplicate marking, position sorting and BAM file indexing were also conducted during the SpeedSeq align procedure using SAMBLASTER(Faust and Hall 2014) and Sambamba (https://github.com/lomereiter/sambamba). The sorted and indexed BAM files were directly loaded into the SpeedSeq somatic module. SpeedSeq analysis using FreeBayes (http://arxiv.org/abs/1207.3907) was performed on the tumor/normal pair of BAM files to achieve somatic SNVs and small INDELs. For the indexed BAM files, we also conducted INDEL realignment and base quality score recalibration using the Genome Analysis Toolkit (GATK)(McKenna et al. 2010), and then the MuTect algorithm(Cibulskis et al. 2013) was applied to identify somatic SNVs. BEDtools was used to filter the somatic SNVs identified by both SpeedSeq and Mutect. Finally, we used the Ensembl Variant Effect Predictor (VEP)(McLaren et al. 2010) to annotate the merged vcf file.

### 4. Pathway analysis

We performed a pathway analysis of the genes that mapped to the somatic SNVs in the tumor samples using DAVID (https://david.ncifcrf.gov/). Genes listed on the DAVID web site and all known human genes were used as the background. A KEGG database analysis was used to evaluate the pathway enrichment.

### 5. Structural variation analysis

LUMPY was run in the SpeedSeq sv module to identify somatic structural variations(Layer et al. 2014), and the vcf files generated by SpeedSeq were annotated using the VEP algorithm. We predicted the somatic structural variations in the primary and lymphatic metastatic GB-SCNECs using DELLY. We identified large somatic deletions, duplications and inversions detected by both LUMPY and DELLY. With respect to the genomic translocational structural variants, we focused on putative gene fusions. Somatic genomic translocational events were generated and filtered by DELLY. OncoFuse(Shugay et al. 2013) was used to identify the putative activating gene fusions.

### 6. Copy number variation analysis

Saas-CNV(Zhang and Hao 2015) was used to study somatic copy number variations in the primary and metastatic GB-SCNEC tissues. The GATK variant analysis pipeline was used to process the data generated from the sequencing system as described above. High-quality SNVs and small INDELs called by GATK were saved in variant call format (VCF) files, and the files were loaded into the Saas-CNV algorithm. The read depth and alternative allele frequency at each site were determined and used to identify somatic copy number alterations (SCNAs).

### 7. Microsatellite stability analysis

We used the vcf2MSAT (http://sourceforge.net/projects/vcf2msat/) script to retrieve the short tandem repeat (STR) data from the vcf files generated from the SpeedSeq somatic module. Information regarding the microsatellite repeat regions identified as INDELs was saved in the vcf files. The data obtained from identifying the genomic location and somatic changes in repeat number were used for the follow-up analyses.

## Results

### 1. Whole-genome sequencing (WGS) and identification of SNVs

We performed WGS of the primary cancer, lymphatic metastatic cancer and matched non-cancerous tissue specimens. The sequencing data were mapped to the human reference genome hg19 with a mean sequencing depth of 61.98X (range 35.73-76.29X) and a mean coverage of 98.7%. Additional information regarding the WGS data is summarized in **Supplementary Table 1**.

Using the SpeedSeq framework and GATK-MUTECT with the non-cancerous gallbladder tissue as the normal reference, we identified 295 and 712 high-quality somatic SNVs in the primary and lymphatic metastatic tumor tissues, respectively (**Figure 2**, **3A**, **3B**). One hundred nine somatic SNVs were shared by the primary and metastatic tumors. Additional information regarding the 109 somatic SNVs is listed in **Supplementary Table 2**. The majority of the identified SNVs were located in intronic and intergenic regions (**Figure 2**). A pathway analysis of the genes mapped to the somatic SNVs revealed a significant enrichment in genes associated with axon guidance, ERBB signaling, sulfur metabolism and calcium signaling (**Figure 4A**). Notably, we identified somatic SNVs in 7 genes (PAK7, HRAS, ERBB4, SOS1, MTOR, NRG1 and ABL1) in the ERBB signaling pathway, which is consistent with the results of our previous study(Li et al. 2014).

**Figure 2.**
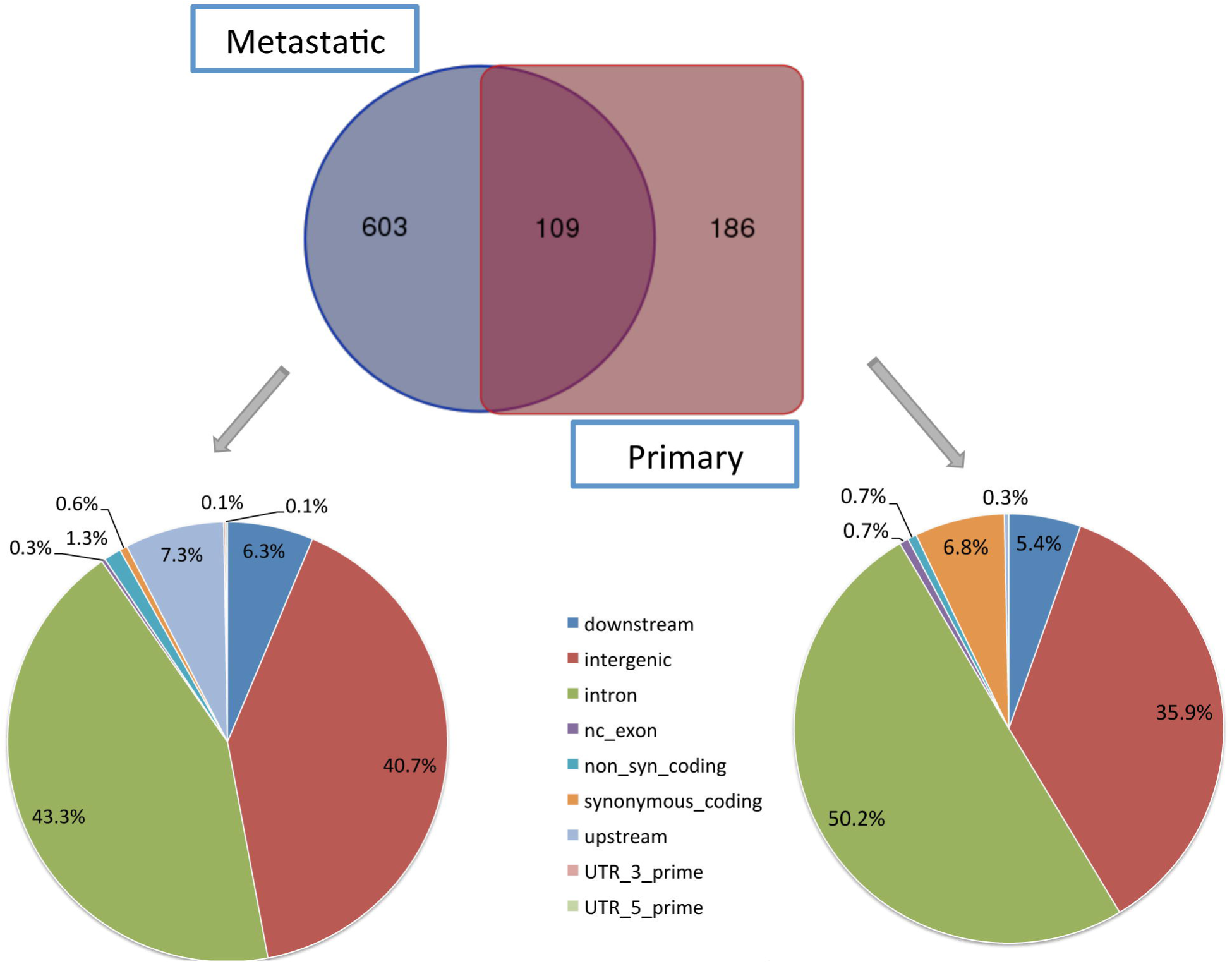
Genomic distribution of SNVs detected in the primary and metastatic GB-SCNEC tissues. In total, 295 and 712 somatic SNVs in primary and lymphatic metastatic tumors were identified, respectively, and 109 somatic SNVs were observed in both the primary and metastatic tumor tissues. The majority of the SNVs were located in intronic and intergenic regions.

**Figure 3.**
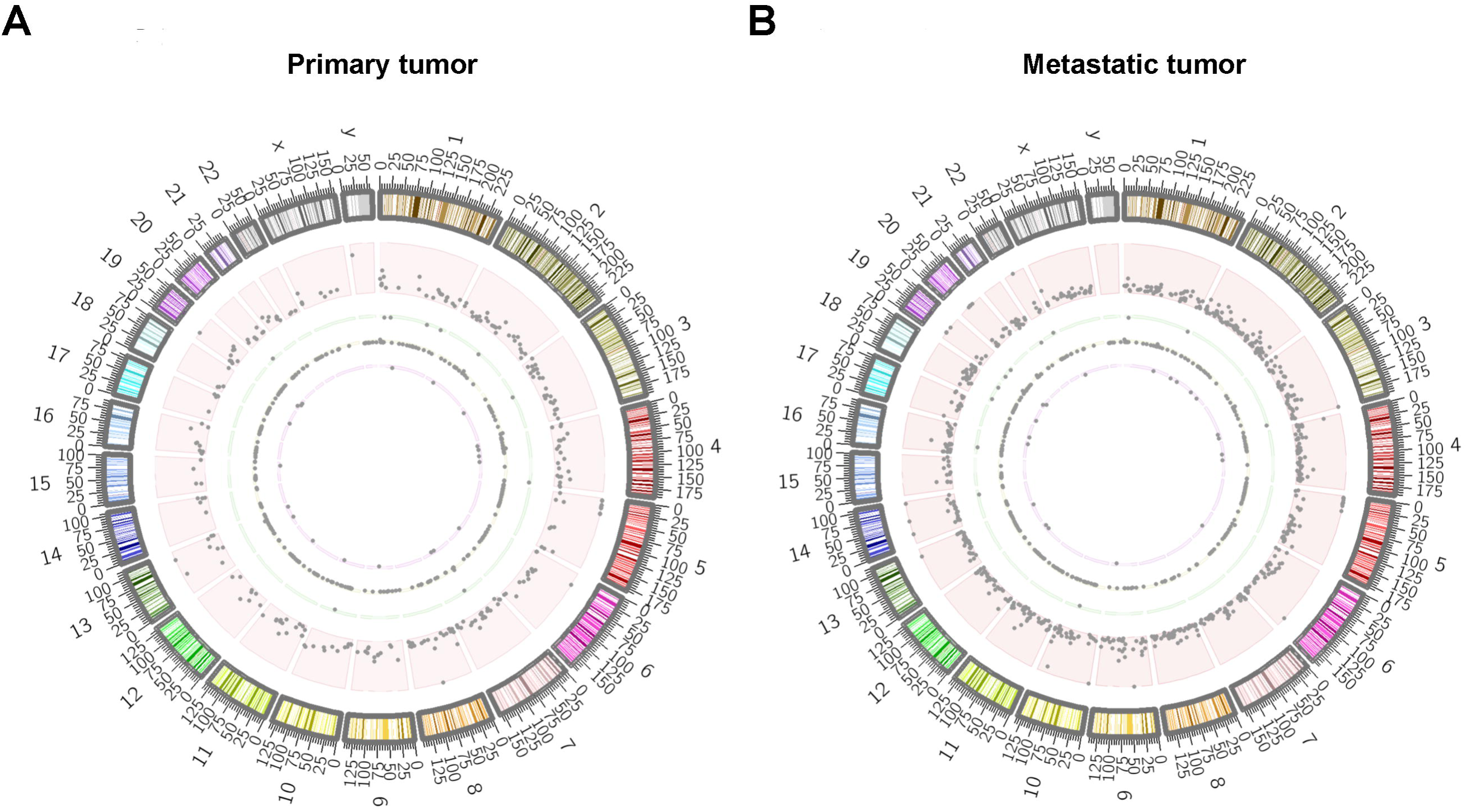
Chromosomal distribution of SNVs, structural variations, copy number variations and microsatellite instability identified in the primary and metastatic GB-SCNECs. (A) Chromosomal distributions of SNVs (red), structural variations (green), copy number variations (yellow) and microsatellite instabilities (purple) observed in the primary GB-SCNEC. (B) Chromosomal distribution of the corresponding variations in the metastatic GB-SCNEC.

**Figure 4.**
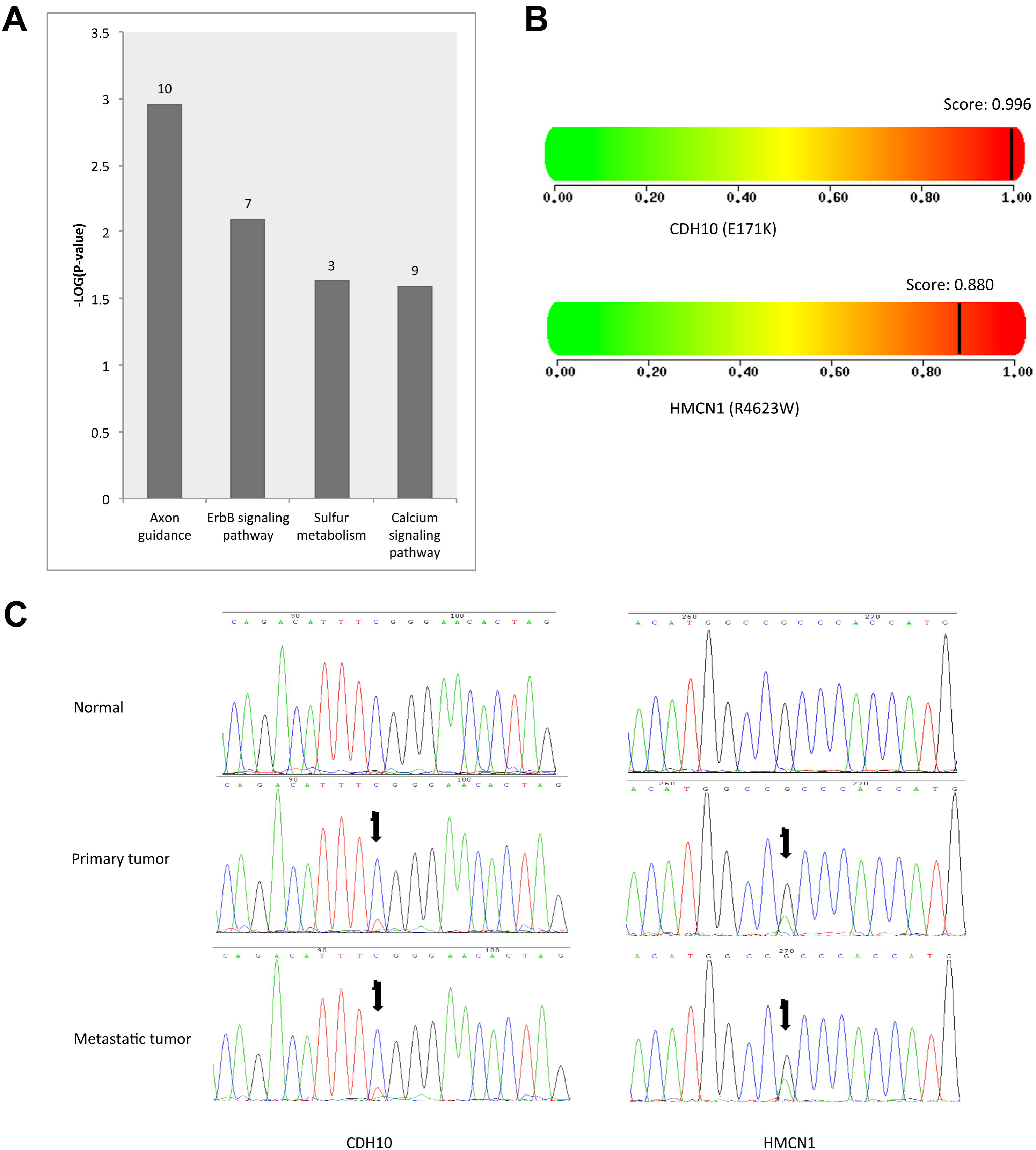
Pathway analysis of the genes with somatic SNVs, functional predictions and sanger sequencing validations of SNVs in HMCN1 and CDH10. (A) Pathway analysis of the somatic genes mapped to SNVs. The genes associated with axon guidance, ERBB signaling, sulfur metabolism and calcium signaling were significantly enriched. (B) Amino acid changes in the proteins encoded by HMCN1 (E171K) and CDH10 (R4623W) had a damaging effect on the structure and function of proteins. (C) The SNVs associated with the amino acid changes were validated using conventional Sanger sequencing.

Next, we investigated the impact of the genetic variations on amino acid changes. Somatic non-synonymous coding variations in HMCN1 and CDH10 were observed in the both the primary and metastatic tissue specimens (**Table 1**). In addition, somatic non-synonymous coding variations in MAST2, C6orf118, UBR5, OR51F1, SLC22A8, CSPG4 and BAIAP2L2 were observed in the metastatic GB-SCNEC tissue. The SNVs associated with amino acid variations were further validated using conventional Sanger sequencing.

**Table1.** Somatic SNVs discovered by whole genome sequencing

| Gene | Chr. | Position | Allele change | Amino acid change | Gerp score | Cadd score | Impact | Tumors observed |
| --- | --- | --- | --- | --- | --- | --- | --- | --- |
| HMCN1 | 1 | 186107047 | C>T | R>W | 3.47 | 3.29 | non_syn_coding | Both |
| MAST2 | 1 | 46487740 | G>A | R>Q | 5.74 | 6.00 | non_syn_coding | Metastatic |
| CDH10 | 5 | 24537504 | C>T | E>K | 5.72 | 3.72 | non_syn_coding | Both |
| C6orf118 | 6 | 165715716 | G>A | T>M | 0.45 | -0.10 | non_syn_coding | Metastatic |
| UBR5 | 8 | 103306037 | C>T | R>H | 6.05 | 4.06 | non_syn_coding | Metastatic |
| OR51F1 | 11 | 4790457 | C>A | A>S | 3.33 | 1.23 | non_syn_coding | Metastatic |
| SLC22A8 | 11 | 62782405 | C>T | R>H | 3.85 | 2.76 | non_syn_coding | Metastatic |
| CSPG4 | 15 | 75982085 | C>T | E>K | 5.26 | 3.58 | non_syn_coding | Metastatic |
| BAIAP2L2 | 22 | 38494168 | C>T | E>K | 5.53 | 5.10 | non_syn_coding | Metastatic |

We further evaluated the impact of the SNVs in HMCN1 and CDH10 on the structure and function of proteins using Polyphen2. We found that the amino acid changes in the proteins encoded by the HMCN1 (E171K) and CDH10 (R4623W) variants were predicted to disrupt the structure and function of proteins (score=0.996 and 0.880, respectively) (**Figure 4B**). The two variants were also clearly validated by Sanger sequencing (**Figure 4C**).

### 2. Analysis of structural variations

Using LUMPY and Delly, we identified more than 50 somatic genomic rearrangements in the primary and lymphatic metastatic GB-SCNECs tissue samples, which included 11 deletions, 4 tandem duplications and 5 inversions mapped to known genes. Additional information regarding the genes that mapped to structural variations is summarized in **Table 2**, and the genomic locations of the structural variations are indicated in **Figure 3A** and **Figure 3B**. Notably, 2 tandem duplications that encompassed the CLCNKA, CLCNKB, FAM131C, C1DP2 and C1DP3 genes were observed in both the primary and lymphatic metastatic GB-SCNEC samples.

**Table2.**
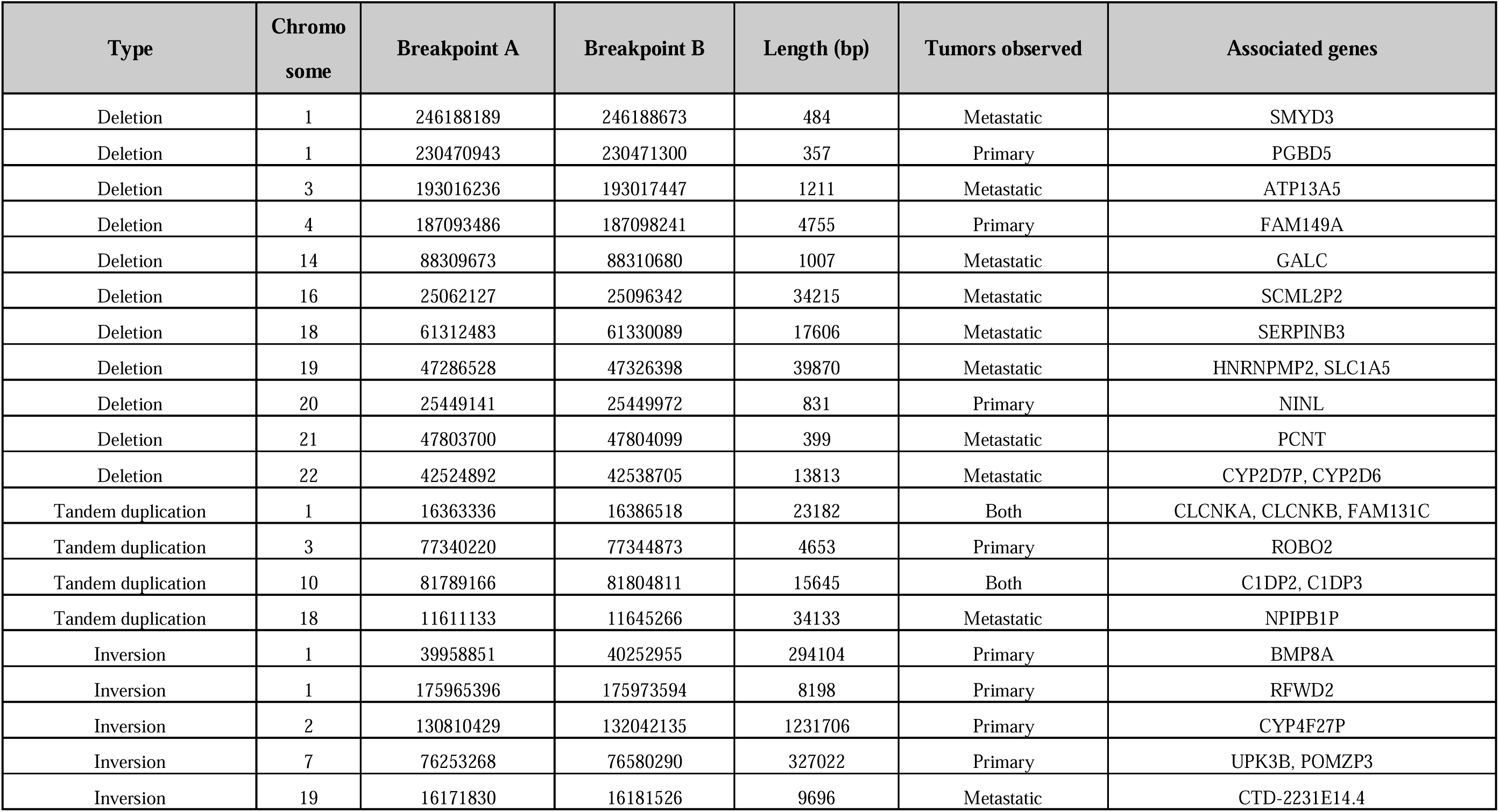
Gene related structure variations discovered by whole genome sequencing

We also considered the tanslocational events generated by DELLY. The high-quality somatic translocational structure variants were filtered. In primary and lymphatic metastatic tumor tissues, we identified 35 and 41 translocations respectively, 10 of which were shared by the primary and metastatic tumors. The detailed information regarding the translocations is summarized in **Supplementary Table 3**. A total of 6 gene fusions were identified, 2 of which, including NCAM2-SGCZ and BTG3-CCDC40 were validated by conventional Sanger sequencing. The genomic locations of all the translocations were labeled in **Figure 5**.

**Figure 5.**
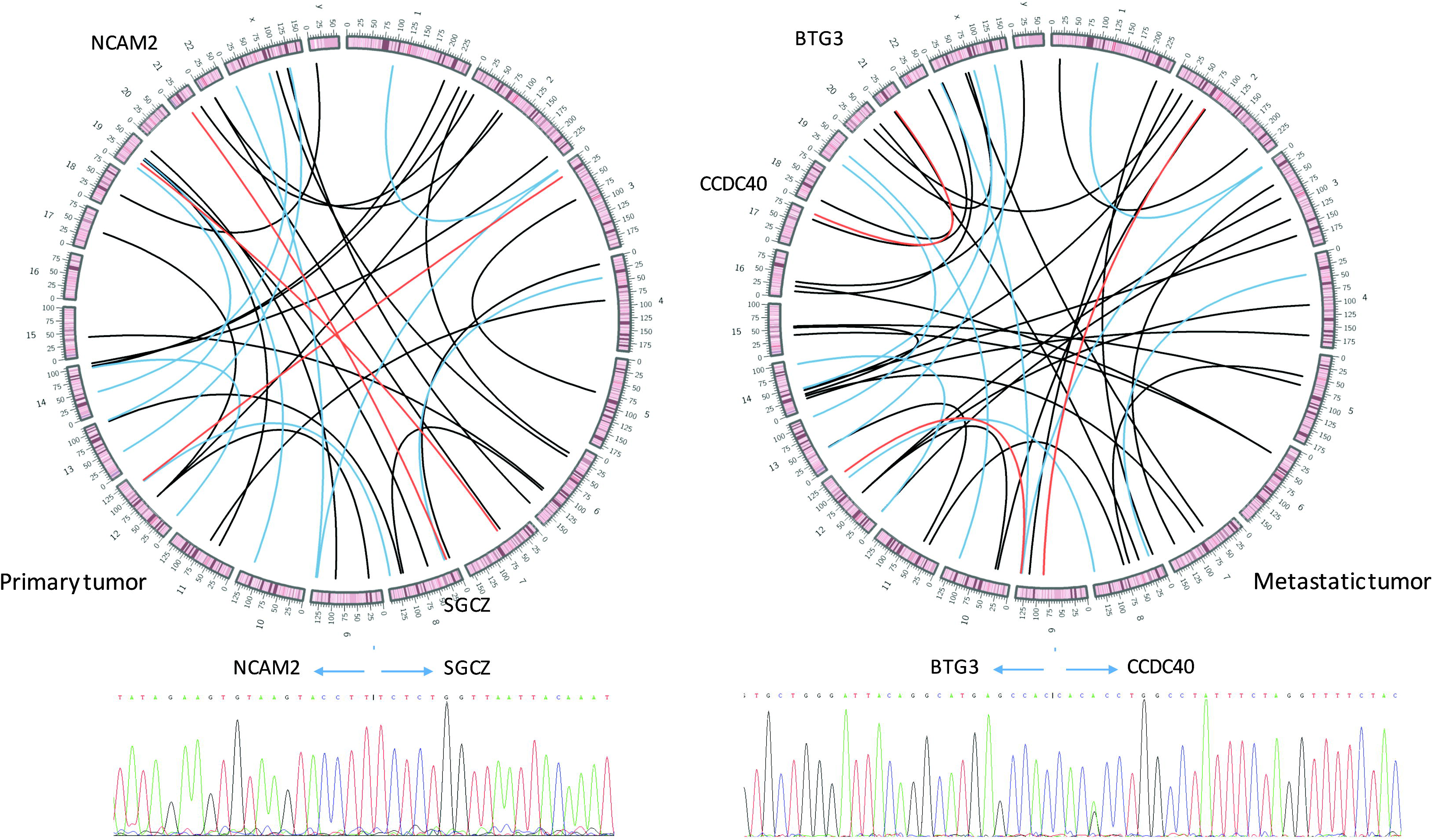
Genomic locations of all the translocational structure variants discovered in the primary and metastatic tumors. The translocatianal regions were linked by lines with different colors (black lines for those specially identified in one of the two tumors, blue lines for those identified in both of the tumors and reds lined for gene fusions).

### 3. Analysis of copy number variants

We identified putative genome-wide copy number variants according to the total read depth and alternative allele proportion at each SNP/INDEL locus. First, we plotted the genome-wide distribution of the alternative allele frequency (also referred to as the B allele frequency or BAF) of the primary and metastatic tumor tissues (**Figure 6**). We found that the general distribution of the BAF was conserved between the 2 primary and metastatic tumor tissues. The SCNAs were estimated by combining the total read depth and the BAF at each locus. A total of 668 SCNAs in the primary tumor and 553 SCNAs in the metastatic tumor were identified. Next, we focused on the SCNAs that mapped to known genes and were observed in both of the primary and metastatic tumors tissues (**Table 3**).

**Table3.** Copy number variations discovered by whole genome sequencing

| Chr | Start | End | Length | Locus number | CNV type | Gene number | Gene name |
| --- | --- | --- | --- | --- | --- | --- | --- |
| chr4 | 3479562 | 3636112 | 156551 | 1912 | loss | 3 | DOK7 et al. |
| chr4 | 34037877 | 45014591 | 10976715 | 8635 | loss | 51 | APBB2 et al. |
| chr5 | 15890 | 3323754 | 3307865 | 4788 | gain | 35 | AHRR et al. |
| chr5 | 21479448 | 21480563 | 1116 | 12 | loss | 1 | GUSBP1 |
| chr5 | 34195054 | 45953915 | 11758862 | 7994 | gain | 66 | AGXT2 et al. |
| chr10 | 49745229 | 51195765 | 1450537 | 894 | loss | 18 | ARHGAP22 et al. |
| chr17 | 171973 | 1755819 | 1583847 | 2072 | loss | 34 | ABR et al. |
| chr17 | 20786423 | 21193432 | 407010 | 438 | loss | 6 | C17orf103 et al. |
| chr17 | 30144873 | 30285495 | 140623 | 51 | loss | 3 | COPRS et al. |
| chr18 | 106728 | 112535 | 5808 | 351 | loss | 1 | ROCK1P1 |
| chr18 | 125075 | 805150 | 680076 | 840 | loss | 9 | C18orf56 et al. |
| chr18 | 23627315 | 24376447 | 749133 | 139 | loss | 5 | KCTD1 et al. |
| chr19 | 8747389 | 9028188 | 280800 | 644 | loss | 5 | ACTL9 et al. |
| chr19 | 17044763 | 20685187 | 3640425 | 2640 | loss | 104 | ABHD8 et al. |
| chr19 | 22618698 | 23481974 | 863277 | 1333 | loss | 6 | LOC440518 et al. |
| chr19 | 34885372 | 36747800 | 1862429 | 1653 | gain | 77 | ALKBH6 et al. |
| chr20 | 25734683 | 25764502 | 29820 | 662 | loss | 1 | FAM182B |
| chr22 | 17278454 | 17338017 | 59564 | 71 | loss | 2 | HSFY1P1 et al. |
| chr22 | 20716839 | 21461982 | 745144 | 964 | loss | 20 | AIFM3 et al. |
| chr22 | 22002021 | 22578991 | 576971 | 930 | loss | 7 | MAPK1 et al. |
| chrY | 2759925 | 9950526 | 7190602 | 2424 | loss | 35 | AMELY et al. |

**Figure 6.**
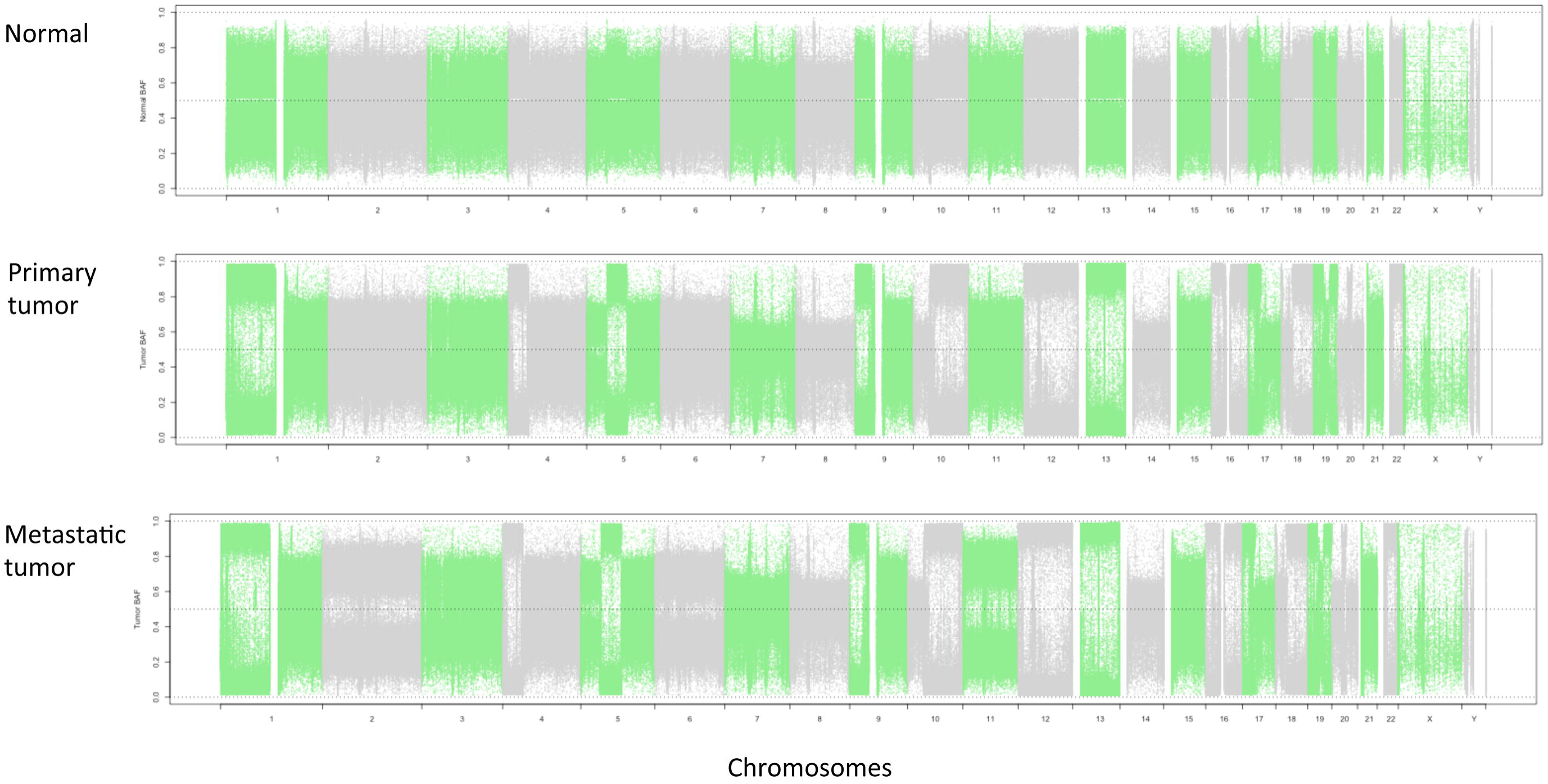
Genome-wide distribution of the alternative allele proportion (also referred to as the B allele frequency, BAF) in the primary and metastatic tumor tissue.

### 4. Microsatellite instability analysis

We extracted the genomic locations of the genomic repeats and the changes in the somatic repeat count identified using the SpeedSeq framework. Additional information regarding the repeat regions identified in the primary and lymphatic metastatic GB-SCNEC tissues is provided in **Supplementary Table 4 and Table 5 respectively**. Repeat motif counts ≥ 4 and somatic count changes > 5 were filtered (**Table 4**). The genomic locations of the repeat motifs with changes in repeat number are also indicated in **Figure 3A** and **Figure 3B**. Genome-wide microsatellite instability was observed in both the primary and metastatic GB-SCNECs tissue, which is consistent with observations from many previous studies.

**Table4.** Microsatellite instability regions discovered by whole genome sequencing

| Chr. | Location | Motif | Length in normal | Length in tumor | Repeat count change | Tumors observed |
| --- | --- | --- | --- | --- | --- | --- |
| chr1 | 69703144 | TAGA | 38 | 66 | 7 | Primary |
| chr1 | 167530083 | GAAA | 91 | 443 | 88 | Metastatic |
| chr2 | 5990493 | TTTA | 74 | 494 | 105 | Metastatic |
| chr2 | 45531195 | TTTC | 45 | 77 | 8 | Both |
| chr2 | 191250792 | TTTTA | 94 | 49 | -9 | Metastatic |
| chr2 | 144313429 | TAGAA | 67 | 27 | -8 | Metastatic |
| chr3 | 28079626 | TTTC | 94 | 362 | 67 | Primary |
| chr3 | 192285524 | TTTC | 123 | 359 | 59 | Metastatic |
| chr3 | 170118525 | TTTC | 124 | 548 | 106 | Metastatic |
| chr4 | 71206459 | TTTA | 19 | 51 | 8 | Both |
| chr4 | 36333809 | TAAAA | 84 | 54 | -6 | Primary |
| chr4 | 13712785 | TAAA | 39 | 67 | 7 | Primary |
| chr4 | 103546995 | TTTTA | 64 | 34 | -6 | Both |
| chr6 | 91122413 | GGAAA | 38 | 68 | 6 | Both |
| chr7 | 92678135 | CAAAA | 54 | 114 | 12 | Primary |
| chr7 | 108898557 | TTTC | 66 | 230 | 41 | Primary |
| chr9 | 74371107 | TAGA | 22 | 46 | 6 | Primary |
| chr9 | 112301512 | TTTC | 113 | 385 | 68 | Metastatic |
| chr10 | 115017138 | TTTC | 123 | 603 | 120 | Primary |
| chr12 | 94909841 | AGAAA | 43 | 88 | 9 | Metastatic |
| chr12 | 19265351 | GAAAA | 71 | 41 | -6 | Primary |
| chr12 | 119158255 | TTTC | 141 | 285 | 36 | Metastatic |
| chr13 | 42832403 | TTTA | 37 | 61 | 6 | Primary |
| chr13 | 104355487 | TTTC | 101 | 73 | -7 | Both |
| chr15 | 84001991 | TTTC | 113 | 577 | 116 | Primary |
| chr16 | 77801681 | GAAA | 51 | 199 | 37 | Metastatic |
| chr17 | 9672167 | TTTA | 36 | 60 | 6 | Metastatic |
| chr18 | 2158902 | GAAA | 75 | 239 | 41 | Primary |
| chr18 | 12298904 | GGGA | 61 | 321 | 65 | Metastatic |
| chr19 | 40429543 | GAAA | 44 | 192 | 37 | Primary |
| chr19 | 31435307 | TAAA | 51 | 107 | 14 | Primary |
| chr19 | 28323640 | TTTTA | 106 | 341 | 47 | Metastatic |
| chr19 | 15971928 | TTTC | 55 | 519 | 116 | Both |
| chrX | 22515660 | TCTA | 62 | 2 | -15 | Metastatic |

## Discussion

GB-SCNEC is a relatively rare and aggressive tumor that is associated with a poor prognosis, and early diagnosis with prompt surgical intervention is currently the only treatment that provides favorable long-term outcomes. Therefore, novel insights into the molecular mechanisms underlying GB-SCNEC are required for the development of effective treatment strategies. In the present study, we performed whole-genome sequencing of primary gallbladder cancer, lymphatic metastatic cancer and matched non-cancerous tissues and identified 295 and 712 somatic SNVs in the primary and metastatic tumor tissues, respectively. The genetic aberrations were mapped to genes associated with axon guidance, ERBB signaling, sulfur metabolism and calcium signaling. Among these genes, HMCN1 and CDH10 were determined to be novel suppressors of gallbladder cancer metastasis. In addition, we identified 11 deletions, 4 tandem duplications and 5 inversions that mapped to known genes. Copy number variants were also analyzed, and genome-wide microsatellite instability was observed in both the primary and metastatic tumor specimens.

We previously reported that the ERBB signaling pathway is associated with the development of gallbladder cancer. Using whole-exome and targeted sequencing technology, we identified 12 genes that are involved in the ERBB signaling pathway and associated with somatic mutations in gallbladder cancer, and they included ERBB4, HRAS and NRG1(Li et al. 2014). In the present study, we identified somatic mutations in ERBB4, HRAS and NRG1 in gallbladder cancer tissue. Together, these findings further validate the critical role of the ERBB signaling pathway in gallbladder carcinogenesis.

CDH10 encodes a type II classical cadherin that mediates calcium-dependent cell-cell adhesion. In this study, we identified a somatic mutation in CDH10 with a deleterious effect on the structure and function of certain proteins. The mutation was observed in both the primary and metastatic tumor tissue, indicating that it is associated with tumor metastasis. A relationship between CDH10 and tumor metastasis has also been observed in colorectal cancer. Jun Yu et al. found that mutations in CDH10 as well as 4 other genes (COL6A3, SMAD4, TMEM132D, VCAN) were significantly associated with improved overall survival in colorectal cancer independent of tumor node metastasis (TNM) stage(Yu et al. 2015). Cadherin-10, the protein encoded by the CDH10 gene, can directly interact with β-catenin, a protein that mediates cell-cell interactions(Shimoyama et al. 2000). Notably, cadherins are calcium-dependent cell adhesion proteins, and we identified mutations in 9 genes involved in calcium signaling.

We also identified mutations in 10 genes that play a role in axon guidance signaling; therefore, this is the first study to report a link between axon guidance signaling and gallbladder neuroendocrine carcinoma. A relationship between axon guidance signaling and neuroendocrine carcinoma has also been observed in small intestine neuroendocrine tumors (SI-NETs) as reported by Banck MS et al.(Banck et al. 2013). Using exome sequencing, these authors found that mutations in axon guidance were frequently disrupted in SI-NET. In summary, disruptions in axon guidance signaling might play crucial roles in neuroendocrine carcinogenesis.

Structural variations that mapped to known genes were also observed in gallbladder neuroendocrine carcinoma tissues. This study is also the first to report the identification of 11 deletions, 4 tandem duplications and 5 inversions in gallbladder cancer. Notably, a 23,182 bp tandem duplication in chromosome 1 and a 15,645 bp tandem duplication in chromosome 10 were observed in both the primary and metastatic tumor tissues. These duplications might disrupt the expression of genes mapped to these regions (CLCNKA, CLCNKB, FAM131C, C1DP2 and C1DP3), and these potential disruptions in gene expression might play a role in tumor malignancy. NCAM2-SGCZ and BTG3-CCDC40 fusions were among the translocational structure variations. Notably, the functional domains in the 4 genes were partially retained in the fusions, which indicated that novel fusion transcripts were involved in the development and metastasis of gallbladder cancer.

This study is also the first to report the identification of copy number variations in primary and metastatic GB-SCNEC tissues. In addition, microsatellite instability has long been considered a contributor to gallbladder carcinogenesis. The genome-wide microsatellite instability observed in the primary and metastatic gallbladder tumor tissues in this study is consistent with the results from many previous reports(Yoshida et al. 2000; Sessa et al. 2003). In summary, this is first study to report the whole-genome sequencing of matched primary and metastatic gallbladder tumor tissues. Our findings have the potential for use in the identification of novel approaches for predicting and treating lymphatic metastasis in gallbladder cancer.

## Acknowledgments

This study was supported by the National Natural Science Foundation of China (No. 81172026, 81272402, 81301816, 81172029, 91440203, 81402403, 81502433 and 31501127), the National High Technology Research and Development Program (863 Program) (No. 2012AA022606), the Foundation for Interdisciplinary Research of Shanghai Jiao Tong University (No. YG2011ZD07), the Shanghai Science and Technology Commission Intergovernmental International Cooperation Project (No. 12410705900), the Shanghai Science and Technology Commission Medical Guiding Project (No. 12401905800), the Program for Changjiang Scholars, the Natural Science Research Foundation of Shanghai Jiao Tong University School of Medicine (No. 13XJ10037), the Leading Talent program of Shanghai and Specialized Research Foundation for the PhD Program of Higher Education-Priority Development Field (No. 20130073130014), the Interdisciplinary Program of Shanghai Jiao Tong University (No.14JCRY05), the Shanghai Rising-Star Program (No. 15QA1403100), Top 6 elitist summit of jiangsu Province (No. 2013-WSN-025), and the People’s Livelihood Science and Technology Demonstration Project of Yixing City (No. 2014-80).

